# Characterization of the Effects of *n*-butanol on the cell envelope of *E. coli*

**DOI:** 10.1101/062547

**Authors:** Eugene Fletcher, Teuta Pilizota, Philip R. Davies, Alexander McVey, Chris E. French

## Abstract

Biofuel alcohols have severe consequences on the microbial hosts used in their biosynthesis, which limits the productivity of the bioconversion. The cell envelope is one of the most strongly affected structures, in particular, as the external concentration of biofuels rises during biosynthesis. Damage to the cell envelope can have severe consequences, such as impairment of transport into and out of the cell; however the nature of butanol-induced envelope damage has not been well characterized. In the present study, the effects of *n*-butanol on the cell envelope of *Escherichia coli* were investigated. Using enzyme and fluorescence-based assays, we observed that 1% v/v n-butanol resulted in release of lipopolysaccharides from the outer membrane of *E. coli* and caused ‘leakiness’ in both outer and inner membranes. Higher concentrations of *n*-butanol, within the range of 2% – 10% (v/v), resulted in inner membrane protrusion through the peptidoglycan observed by characteristic blebs. The findings suggest that strategies for rational engineering of butanol-tolerant bacterial strains should take into account all components of the cell envelope.

## INTRODUCTION

Dwindling levels of fossil fuels and the urgency to reduce greenhouse emissions have increased the need for alternative and affordable sources of liquid transportation fuels, which include biofuels. Butanol is a promising next generation biofuel since it has similar properties to gasoline and can be used in gasoline engines without the need for modification (Savage 2011). Its production from fermentation has been demonstrated using native (Green 2011) or engineered bacterial cell factories such as *Escherichia coli* (Bond-Watts et al. 2011). However, it is particularly toxic to microbial cells and has, therefore, struggled to compete with fossil fuels due to low yields and titres obtained from the production (Steen et al. 2010; Wang et al. 2011). In order to prevent product accumulation, which is the main cause of butanol toxicity, several downstream processes such as gas stripping and pervaporation are used to remove the product as it forms, resulting in high downstream processing costs (Van Hecke et al. 2016). An alternative strategy is to increase the host’s tolerance to butanol and, thus, cut down the costs involved in downstream processing. To ensure that this strategy is successful it is important to fully understand the mechanisms that lead to butanol toxicity in the production strain.

The cell envelope of bacteria is a primary point of contact of stresses from the environment including changes in pH, temperature, osmolarity and solvent toxicity (Hommais et al. 2002; Mansilla et al. 2004; Mrozik et al. 2004). Therefore, it is an important target of butanol toxicity as butanol accumulates in the bioreactor during biosynthesis and tries to re-enter the cell. In gram-negative bacteria such as *E. coli*, the cell envelope is composed of an outer and inner membrane with a stiffer peptidoglycan layer between (Ruiz et al. 2006). The outer membrane is asymmetrical and made of phospholipids in the inner leaflet and mainly lipopolysaccharides (LPS) on the outer leaflet (Strain et al. 1983). The LPS comprises three components – the lipid A that anchors the LPS into the hydrophobic region of the outer membrane, a core oligosaccharide containing 2-keto-3-deoxyoctonic acid (KDO) and an O-antigen attached to the oligosaccharide (Raetz and Whitfield 2002). Newly synthesised LPS molecules are transported to the outer membrane in a process chiefly mediated by the LptA protein encoded by the *lptA* gene (Chng et al. 2010). Previous work found that the *lptA* gene is upregulated when *E. coli* is exposed to toxic concentrations of butanol, suggesting that butanol causes membrane stress through damage to the LPS (Reyes et al. 2012). However, the exact mechanism as well as whether the damage to the LPS is the main and/or only butanol induced damage is not known.

Here, we investigate the effect of extracellular *n*-butanol on key components of the cell envelope using *E. coli* as the model organism. Briefly, by developing inner and outer membrane damage reporters we show that damage to the *E. coli* cell envelope is complex and affects both the inner and outer membranes and at higher concentrations the function of the cell wall. Our work, ultimately, aims to guide the development of new strategies required for strengthening the cell envelope of bacterial hosts exposed to *n*-butanol.

## MATERIAL AND METHODS

### Bacterial strains, plasmids and growth conditions

*E. coli* MG1655 (*F*^-^ *λ*^-^ *ilvG*^-^ *rfb-50 rph-1*) was used in this study for all experiments described. The strain was transformed with the following plasmids: (a) for the purpose of studying inner and outer membrane damage we constructed the pIMD and pOMD plasmids respectively, (b) for the bioreporter assays we constructed pBioReporter plasmid; (c) for the purpose of microscopy studies of cell shape changes we used pWR20 and (d) for the purpose of studying SOS response induction we used p15 (Zaslaver et al. 2006). Additionally, for the bioreporter assay we used MG1655 with a chromosomal insertion of Yellow Fluorescence Protein (MG1655-YFP) (Elowitz et al. 2002) as a control. All the plasmids used in the study are listed in Table S1 of the Supplementary Materials. pOMD, pIMD and pBioReporter are derivatives of pSB1C3 obtained from the Registry of Standard Biological Parts (Registry of Biological parts 2008) carrying the ColE1 *ori* and a chloramphenicol resistance marker. pBioReporter was constructed in the following way: The *lptA* promoter was amplified by PCR from the *E. coli* MG1655 genome and was standardised into the BioBrick format (Knight 2003). The PCR was carried out with the high fidelity Phusion DNA polymerase from New England Biolabs Inc. (NEB) using the following pair of primers:

Forward: ATC<u>GAATTC</u>CT<u>TCTAGA</u>GATAACGCGCAGATCAATCTGGTGACGC

Reverse: ATC<u>CTGCAG</u>CT<u>ACTAGT</u>AGGATGTTCTAACCTTTTCAATCAGCTCGGCG

The forward primer was designed to include the BioBrick prefix while the reverse primer included the BioBrick suffix sequence. The primers were synthesised by Sigma-Aldrich. The *lptA* promoter was placed upstream of a ribosome binding site (RBS) and a red fluorescent protein (RFP). Both RBS and RFP were obtained from the Registry of Standard Biological Parts (Registry of Biological parts 2008) as BBa_B0034 and BBa_E1010 respectively. The *lptA* promoter + RFP construct was *EcoRI/PstI* digested and ligated into pSB1C3. The insertion was sequence verified using vector-specific sequencing primers, pSBNX3 insf2 (AAATAGGCGTATCACGAGGC) and pSBNX3 insr2 (CAGTGAGCGAGGAAGCCTGC). pWR20 plasmid carries a gene encoding an enhanced green fluorescent protein (EGFP) as well as a kanamycin resistance gene (Pilizota and Shaevitz 2012). EGFP constitutively expressed from the plasmid freely diffuses in the cytoplasm and thus marks the cytoplasmic volume. p15 is derived from pUA66, which carries pSC101 origin and kanamycin resistance gene, is of relatively low copy number and carries a *recA* promoter controlling the expression of mGFP_mut2 fluorescent protein (Zaslaver et al. 2006).

*E. coli* was aerobically grown in Lysogeny broth (LB), on solid LB agar or in M9 medium (4xM9 salts (28 g/l NaHPO4, 12 g/l KH_2_PO_4_, 2 g/l NaCl, 4 g/l NH_4_Cl), 10 mg/ml thiamine hydrochloride, 0.4% glucose and 0.2% casamino acid). Media were supplemented with the appropriate antibiotics for selection during growth and all of the growth was performed at 37°C with shaking.

### Enzyme assays

Samples used for alkaline phosphatase assay were prepared as follows: an overnight culture of MG1655 carrying pOMD (Table S1) was re-grown in LB (following 1:100 dilution in fresh medium) supplemented with 90 μg/ml IPTG until attaining an OD_600_ of 0.5-0.6. The liquid culture was then split into four and 1% v/v *n*-butanol, 8 mM EDTA, 8 mM SDS or water (no solvent control) was added to each. The cultures were further incubated for 1 h at 37°C and were centrifuged afterwards at 13000 g for 3 min to obtain the supernatants which were used for the assay.

Alkaline phosphatase activity was determined as follows: The reaction mixture contained 0.05 M Tris-HCl (pH 8.0), 2 mM MgCl_2_, 50 mM NaCl, 20 mM *p*-nitrophenyl phosphate (*p*-NPP) substrate and 1:10 dilution of sample (supernatant) prepared as described above. To determine the effect of 8 mM EDTA on the outer membrane, a higher concentration (40 mM) of MgCl_2_ was used to prevent EDTA from inhibiting the alkaline phosphatase. The reaction mixture was incubated at 37°C until a yellow product (*p*-nitrophenol) was formed. The time taken for the mixture to turn yellow was noted as incubation time (min). The reaction was stopped by adding 1 M potassium phosphate (KH_2_PO_4_) to the final concentration of 400 mM. Enzyme activity was determined as:

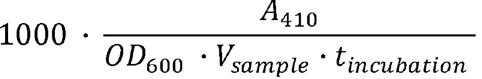

where A_410_ is the absorbance of the reaction mixture measured at 410 nm, OD_600_ the optical density at 600 nm, and V_sample_ and t_incubation_ volume of the sample and the incubation time respectively.

### Fluorescence assay to detect inner membrane damage

*E. coli* MG1655 carrying pIMD (Table S1) was grown in M9 medium until log phase (OD_600_ 0.5-0.6) and then exposed to different concentrations of *n*-butanol (0%, 0.5%, 0.7%, 0.9%, 1.1% v/v). The cultures were then incubated for 1 h at 37 °C after which they were centrifuged (13000 g for 3 min) to separate the cells from the liquid medium. Aliquots of the supernatants were placed in cuvettes and fluorescence measurements were taken with the Modulus Multimode reader, green filter (Turner Biosystems).

### Bioreporter assay

MG1655-YFP expressing RFP from the *lptA* promoter on pBioReporter (Table S1) was grown in M9 from an overnight culture to OD~0.9 (log phase) after which different concentrations of *n*-butanol or EDTA were added to cultures. Red and yellow fluorescence were measured at different time points. The data was normalized by dividing the RFP output from the bioreporter by the YFP signal (yellow fluorescence) produced constitutively by the MG1655-YFP strain.

### Lipopolysaccharide (LPS) assay

Overnight culture of MG1655 strain was regrown in LB (following 1:100 dilution in fresh medium) to OD_600_ ~0.7. The cells were harvested by centrifugation, washed three times and then re-suspended in 1×PBS (137 mM NaCl, 2.7 mM KCl, 10 mM Na_2_HPO_4_ and 1.8 mM KH_2_PO_4_) to remove any residual growth medium. *n*-Butanol or EDTA was added to cell suspensions and incubated with shaking for 1 h. The cell suspensions were centrifuged at 13000 g for 3 min to obtain the supernatant containing released LPS for the assay.

The purpald assay was used to quantify LPS released from the outer membrane as described by Lee & Frasch (2001) with slight modifications. All the reagents used for the assay were freshly prepared before use. The supernatants obtained as described above were treated with 32 mM sodium periodate (NaIO_4_) and incubated for 25 min. Then, 136 mM purpald reagent was added and incubated for 20 min followed by the addition of 64 mM NaIO to the reaction mixture. After incubating for another 20 min, the absorbance was measured at 550 nm.

### Microscopy

Overnight culture of MG1655 with pWR20 and MG1655 with p15 were regrown (1 in 100 dilution) in LB supplemented with different concentrations of *n*-butanol (0%, 0.5% and 1% v/v) to OD_600_ ~0.5. To investigate the effect of harsher concentrations on the cell, MG1655 with pWR20 was grown in LB to OD_600_ of 0.5 after which n-butanol was added to different aliquots of the culture to a final concentration of 1.5%, 2%, 2.5%, 3%, 4% and 5% v/v and incubated for 15 min. Samples were prepared by placing a drop of cell culture between the microscope slide and the cover slip. The cells were allowed to settle to the glass surface for 5 min after which the sample was observed in epifluorescence using the Zeiss Axiovert 200 fluorescence microscope equipped with a photometrics cool-SNAP HQ CCD camera.

## RESULTS

### Detection of outer membrane damage

In order to determine the effect of n-butanol on the outer membrane of *E. coli*, leakage of the periplasmic protein, alkaline phosphatase (PhoA), was measured. In the exponential growth phase PhoA is not expressed and levels present are negligible (Heppel et al. 1962). Thus, we overexpressed the *phoA* gene in the pOMD plasmid (Table S1) and evaluated the effect of *n*-butanol on the alkaline phosphatase activity to ensure that the enzyme disruption due to butanol toxicity does not occur during the assay. Indeed, we found that the enzyme activity was not altered in the presence of n-butanol (Fig. S1). Next, we assayed the supernatants of cultures exposed to 1% v/v *n*-butanol, 8 mM EDTA and 8 mM SDS for 1 h to detect alkaline phosphatase activity. EDTA and SDS were included as positive controls, as they are known to have negative effects on the cell membranes (Woldringh and Van Iterson 1972; Hardaway and Buller 1979). EDTA targets the outer membrane whereas SDS dissolves both inner and outer membranes. We detected alkaline phosphatase activity in the supernatants of these cultures with the *n*-butanol cultures having a 2-fold higher alkaline phosphate activity than the control cultures which contained neither *n*-butanol, EDTA nor SDS (Fig. 1). The positive control cultures, EDTA and SDS had a 1.4 and 8 fold enzyme activity higher than control cultures respectively (Fig. 1). SDS had the strongest effect on the cell membrane which was expected since SDS is known to solubilise cell membranes (Singer and Tjeerdema 1993). We also expected the cultures treated with EDTA, which has been reported to permeabilize bacterial membranes (Hardaway and Buller 1979), to show high enzyme activity in the supernatant, but we observed a rather low activity. This was true even when excess amounts (4 times as much EDTA) of Mg^2+^ was added to the supernatants used for the assay to prevent EDTA from inhibiting the alkaline phosphatase.

**Fig. 1.**
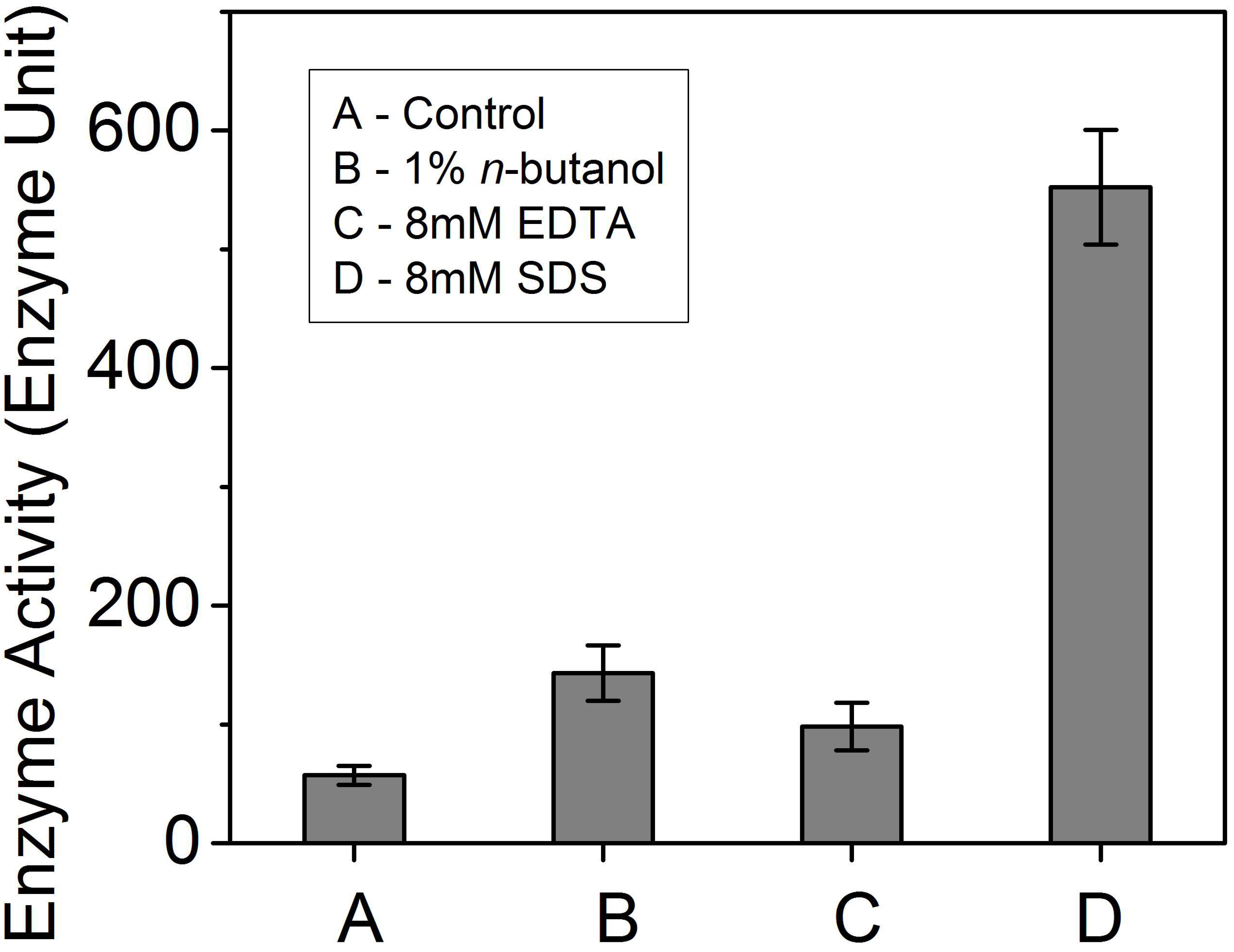
Effect of *n*-butanol on the outer membrane of *E. coli* as measured by alkaline phosphatase activity. PhoA leakage from the periplasm into the supernatant of cultures exposed to n-butanol was used to assess the effect of n-butanol on the outer membrane. EDTA and SDS were used as controls. Experiments were done in triplicate. Error bars indicate one standard error.

To confirm our results obtained with the alkaline phosphatase assay, we developed an additional outer membrane damage assay. The *lptA* bioreporter assay was used to detect *n*-butanol-induced outer membrane damage as described in Materials and Methods. We chose the *lptA* promoter because it is upregulated during cell stress as a result of membrane damage requiring the need for LPS synthesis (Sperandeo et al., 2007 and Reyes et al., 2012). The red fluorescent protein encoded on pBioReporter was used as a quantitative marker of *lptA* induction indicating outer membrane damage. In order to eliminate the effects of the reduced growth rate when *n*-butanol is present (Fig. S2) we used cells that were constitutively expressing YFP (MG1655-YFP) to normalize the bioreporter output. The bioreporter responded to the presence of n-butanol in the medium with a corresponding increase in output over time (Fig. 2A). We also observed a further increase in output when the concentration of n-butanol was increased. To our surprise, the bioreporter was not responsive to the presence of EDTA (control) in the medium even when the concentration of EDTA was increased (Fig. 2B).

**Fig. 2.**
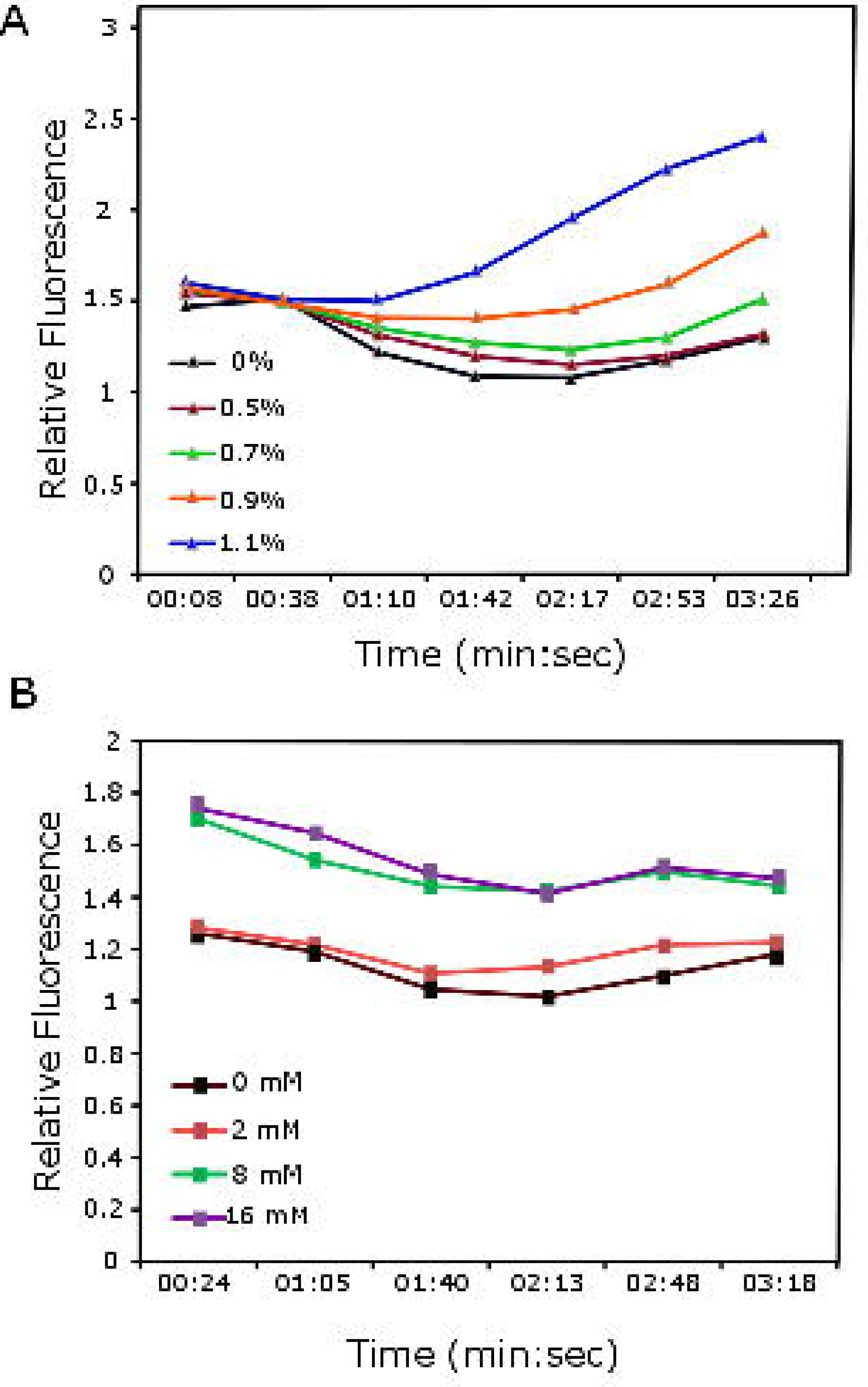
Characterization of the *lptA* bioreporter. The graphs show the effect of increasing concentrations of *n*-butanol (A) and EDTA (B) on the induction of the *lptA* bioreporter. The relative fluorescence was determined as a measure of red fluorescence (from the bioreporter) to a measure of constitutively expressed YFP in the MG1655-YFP. Experiments were done in triplicate.

### LPS release from the outer membrane

To test the hypothesis that *n*-butanol damages the outer membrane by removing the lipopolysaccharides, we grew cells until they reached the early exponential phase and, subsequently, treated with *n*-butanol for 1 h. We observed an increase in the amount of LPS released into the supernatant with increasing concentrations of *n*-butanol (Fig. 3). A similar trend was observed when cells were treated with increasing concentrations of EDTA with EDTA having a stronger effect than *n*-butanol (Fig. 3).

**Fig. 3.**
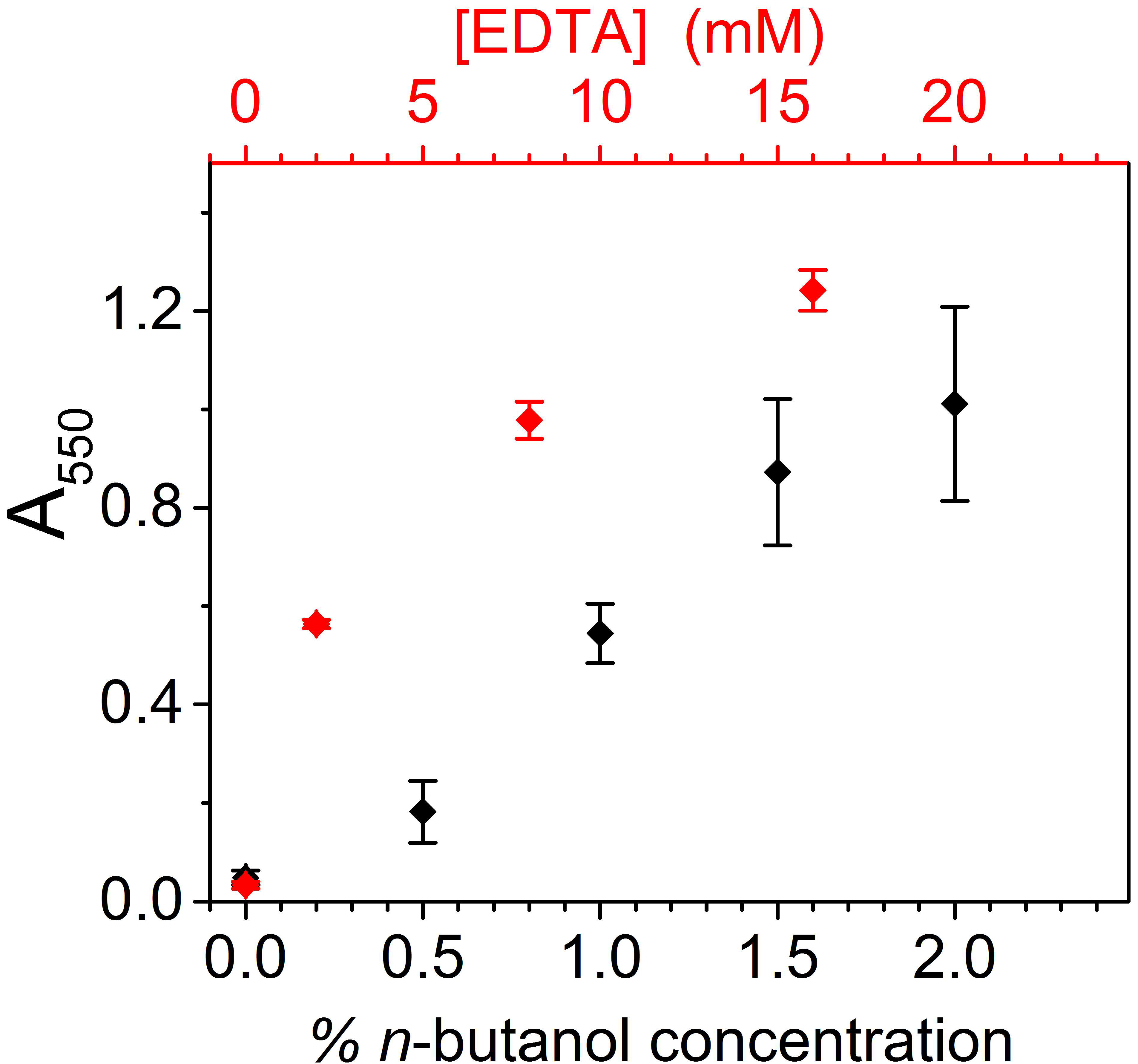
Effect of *n*-butanol on the LPS. Removal of LPS from the outer membrane into the supernatant by *n*-butanol was determined with the purpald assay (shown in black). EDTA was used in a control experiment (shown in red). Experiments were done in triplicate. Error bars indicate one standard error.

### Detection of inner membrane damage

Next, we explored the possibility of *n*-butanol causing damage to the inner membrane too, by measuring the presence of RFP (DsRed), with a molecular size of 28 kDa (Baird et al. 2000), in the supernatants of cultures exposed to increasing concentrations of *n*-butanol (Fig. 4). These cultures were established from *E. coli* MG1655 with pIMD plasmid (carrying the *rfp* gene under the control of a strong promoter (*lac*) see also Materials and Methods and Table S1). We detected a corresponding increase in red fluorescence in the supernatants of these cultures exposed to increasing concentrations of *n*-butanol suggesting RFP leakage from the cytoplasm which is indicative of inner membrane damage.

**Fig. 4.**
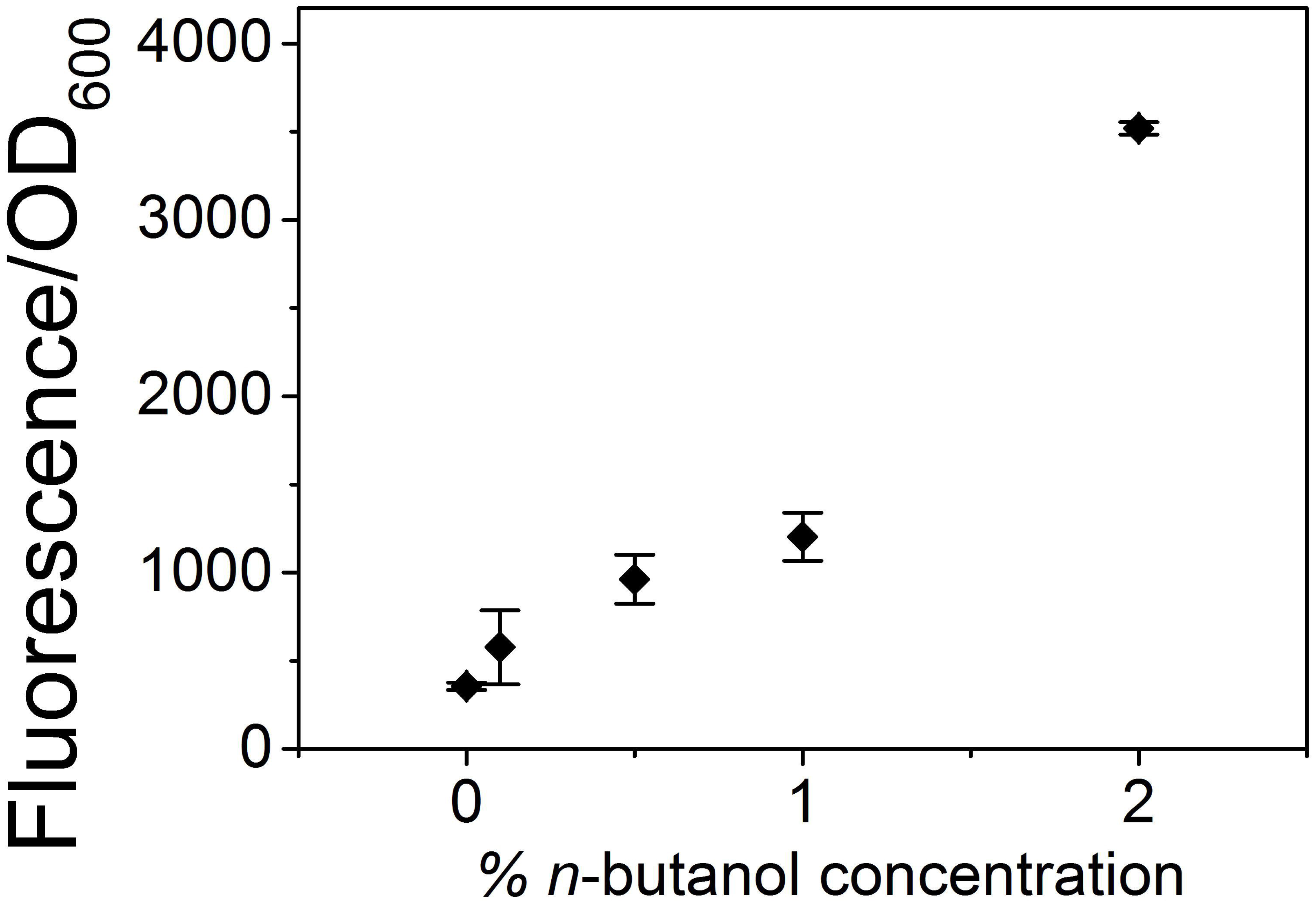
Effect of *n*-butanol on the inner membranes of *E. coli*. The graph shows the effect of increasing concentrations of *n*-butanol on the inner membrane of *E. coli*. Red fluorescence was measured in the supernatants of liquid cultures to determine cytoplasmic leakage. Experiments were done in triplicate. Error bars indicate one standard error.

### Detection of peptidoglycan damage

To determine the effect of *n*-butanol on the cell wall of *E. coli*, we grew MG1655 with pWR20, expressing EGFP and freely diffusing in the cytoplasm, in LB supplemented with 0%, 0.5% and 1% v/v *n*-butanol. To test the limits of *n*-butanol toxicity we investigated even higher concentrations of *n*-butanol, 1.5%, 2%, 2.5%, 3%, 4% and 5% (v/v) as described in *Materials and Methods*. The cells in the control cultures (0% v/v *n*-butanol) and those containing 0.5% v/v *n*-butanol maintained their rod-like shape (Fig. 5A and 5B). However, those exposed to 1% v/v *n*-butanol were elongated and filamentous (Fig. 5C). One possible explanation for this phenotype is DNA damage and induction of SOS response (Bi and Lutkenhaus 1993). To test this possibility, we grew MG1655 with p15 plasmid in LB supplemented with 0%, 0.5% and 1% v/v *n*-butanol. RecA is part of the SOS response and p15 enabled us to monitor the expression of *recA*, and thus SOS response, using a GFP protein marker (see also *Materials and Methods)*. As a positive control we used antibiotic nalidixic acid, which targets DNA and is known to induce SOS damage (Lewin et al. 1989). Fig. S3 shows induction of SOS response in the positive control (A and B) but not in the filamentous cells grown in 1% v/v *n*-butanol (G and H). For the cells exposed to 2%, 3%, 4% and 5% v/v *n*-butanol, we observed the formation of characteristic blebs (Fig. 5D). The cells exposed to 4% and 5% *n*-butanol formed more blebs and at a faster rate than those exposed to lower *n*-butanol concentrations (Figs. 5E, S3). We observed fluorescence in the blebs indicating that they are filled with cytoplasmic contents of the cell (eg EGFP) that protrudes through the cell wall. Furthermore, at the concentrations at which severe morphological changes are observed growth is no longer supported (Fig. 5F).

**Fig. 5.**
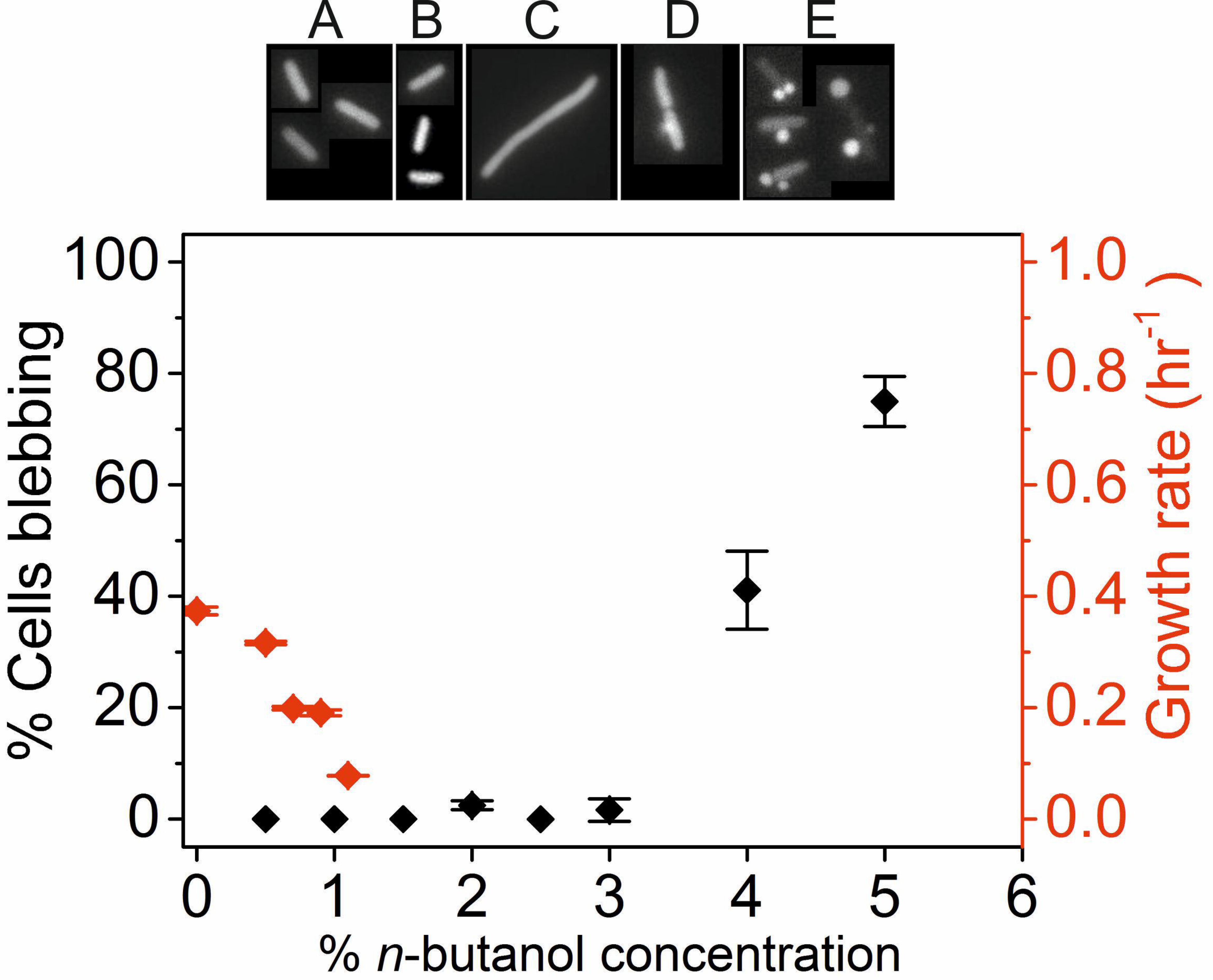
Effect of *n*-butanol on the morphology of *E. coli*. (A-E) Images show changes in cell shape upon exposure to increasing concentrations of *n*-butanol. Shown in A are cells grown with no *n*-butanol present, in B and C are cells grown in the presence of 0.5% and 1% v/v *n*-butanol respectively. Cells in D and E were grown to mid-exponential phase before being exposed to 2% and 5% *n*-butanol respectively. The graph shows the percentage of cells that formed blebs (in red) and changes in growth rate (in black) of cells upon exposure to increasing concentrations of *n*-butanol. The percentage of cells that form blebs was counted from 50-100 cells in each condition.

## DISCUSSION

While the exact damaging effect of butanol on the cell envelope is not known, butanol has been reported to intercalate into the cell membrane resulting in increased fluidity of the membrane (Ingram 1986). Our aim was to investigate and characterise the effect of *n*-butanol on the integrity of *E. coli* cell envelope further.

We observed that both outer and inner membranes of *E. coli* were compromised in cells exposed to *n*-butanol resulting in leakage of periplasmic and cytosolic proteins. In testing the membrane ‘leakiness’ caused by *n*-butanol, we observed that a high level of membrane damage induced by SDS did not inhibit growth as much as *n*-butanol did even though the latter induced less membrane leakage (Fig. S4). This is in agreement with previous studies (Rutherford et al. 2010; Huffer et al. 2011; Gonzalez-Ramos et al. 2013) and confirms that although weakening of the membrane is detrimental to the cell it is not the sole cause of butanol-induced growth inhibition. An interesting observation made in this study was that EDTA, used as a positive control to detect outer membrane damage showed strong removal of LPS from the outer membrane but did not render the outer membrane as ‘leaky’ as in the case of *n*-butanol. This suggests that the removal of LPS is not the only mechanism by which the outer membrane becomes leaky in the presence of *n*-butanol.

Indeed, most hydrophobic compounds in the extracellular environment of the cell are prevented from traversing the membrane into the cell by the hydrophilic lipopolysaccharides (LPS) (Leive 1974). It has been shown previously that when *Pseudomonas aeruginosa* cells were treated with rhamnolipid to release the outer membrane LPS the cells become permeable to organic solvents such as hexadecane (Al-Tahhan et al. 2000). Since *n*-butanol is hydrophobic, it was hypothesised in this study that for *n*-butanol to reach the lipid bilayer from the outside of the cell, it will have to cross the LPS barrier. Our results show that *n*-butanol compromises the integrity of the outer membrane by removing the LPS which maintains the stability of the membrane. However, our results with EDTA strongly suggest that this is not the only cause of damage to the outer membrane by the presence of *n*-butanol. While outer membrane disruption will potentially make the membrane more permeable to *n*-butanol, *n*-butanol might also cross the outer membrane into the cell by other routes, such as through membrane porins.

Using fluorescence microscopy, we showed that exposure to relatively mild concentrations of *n*-butanol (1% v/v) resulted in cell elongation. Formation of reactive oxygen species has been observed in *n*-butanol stressed *E. coli* (Rutherford et al. 2010), which can induce DNA damage and, subsequently, inhibit cell division. We tested this hypothesis and found no SOS response induction in filamentous cells grown in the presence of 1% v/v *n*-butanol. Instead, filamentation is most likely the result of changes in the cell membrane disrupting the function of the MinDC or FtsZ protein systems directly associated with septum formation (Bi and Lutkenhaus 1993). Exposure to harsher concentrations of *n*-butanol (2% to 5% v/v) resulted in a different kind of response (bleb formation) which is characteristic of cell wall damage (Yao et al. 2012). Bleb formation has also been shown in *E. coli* cells treated with β-lactam antibiotics which are known to inhibit cell wall biogenesis (Yao et al. 2012). The blebs formed on the *E. coli* cells suggest that very high concentrations of *n*-butanol result in damage to the peptidoglycan layer. This effect of *n*-butanol on the peptidoglycan has not been previously reported to the best of our knowledge, although some transcriptional studies have reported the upregulation of genes involved in peptidoglycan biosynthesis during isobutanol stress (Atsumi et al. 2010). The speed at which the bleb formation occurred (28 sec after the butanol shock; see video in Supplementary Materials) may suggest rapid denaturation of proteins involved in peptidoglycan biosynthesis. We do not exclude an alternative hypothesis to bleb formation, which is that they are the consequence of a more fluid inner membrane. However, in both scenarios the function of the cell wall, as a stiffer component in the periplasm, is compromised. Although currently these high concentrations are not sustaining growth, they were used in the study to identify components of the cell that need to be engineered in the future to allow growth at higher *n*-butanol concentrations. This is of interest to commercial production of butanol that depends on the production of high titres of *n*-butanol in order to cut down purification costs. It will be interesting, therefore, to further characterise the effect of *n*-butanol on the peptidoglycan layer as this may be a previously unappreciated mechanism of toxicity.

Taken together, these results indicate that *n*-butanol has severe consequences on the cell by, at least, targeting different components of the cell envelope. Hence, engineering bacterial cells factories for the production of toxic biofuels such as *n*-butanol will require the use of robust strategies to protect the cell envelope as a whole from product-induced damage.

## Compliance with Ethical Standards

### Funding

EF would like to acknowledge the Darwin Trust of Edinburgh for the PhD studentship.

### Conflict of interest

The authors declare that they have no conflict of interest.

### Ethical approval

This article does not contain any studies with human participants or animals performed by any of the authors.

